# Monocyte-Mimetic Nanoparticle Delivery of Verteporfin Promotes Plaque-Stabilizing Remodeling in Atherosclerosis

**DOI:** 10.64898/2026.09.16.750992

**Authors:** Ting-Yun Wang, Andrew Zehnder, Sin-Pei Wu, Hui-Chun Huang, Christopher L. Plaisier, Kuei-Chun Wang

**Affiliations:** School of Biological and Health Systems Engineering, Arizona State University, Tempe, USA; School for Engineering of Matter, Transport and Energy, Arizona State University, Tempe, USA; John Shufeldt School of Medicine and Medical Engineering, Arizona State University, Tempe, USA

**Keywords:** biomimetic nanoparticles, targeted nanotherapy, atherosclerosis, vascular inflammation, plaque remodeling

## Abstract

**Background:** Despite advances in lipid-lowering and anti-inflammatory therapies, pharmacological strategies that act directly within established atherosclerotic lesions to suppress plaque progression remain limited. We recently developed a monocyte-mimetic nanoparticle (MoNP) platform that selectively targets inflamed endothelium and delivers Verteporfin (VP), termed MoNP-VP, to suppress YAP/TAZ-associated endothelial activation and plaque development. Here, we extend this biomimetic nanotherapy by defining the pharmacological effects of MoNP-VP within atherosclerotic vessels and evaluating its therapeutic efficacy in pre-existing plaques under distinct lipid burden.

**Methods:** MoNP-VP were formulated by encapsulating VP in polymeric cores followed by cloaking with mouse monocyte membranes. Partial ligation was performed in ApoE-deficient mice, and single-cell RNA sequencing was used to define pharmacodynamic responses in carotid lesions. Proprotein convertase subtilisin/kexin type 9 (PCSK9) gain-of-function was induced in wild-type mice to generate pre-existing aortic lesions, followed by continued high-fat-diet or switching to chow to evaluate MoNP-VP efficacy under persistent hyperlipidemia and lipid-lowering conditions.

**Results:** Single-cell analysis revealed that MoNP-VP remodeled the plaque microenvironment in carotid arteries, selectively reducing foamy macrophage populations while enriching vascular stromal populations associated with fibrotic remodeling. In mice with pre-existing aortic plaques, MoNP-VP suppressed lesion progression and reduced macrophage content under persistent hyperlipidemia. Following diet-induced lipid lowering, MoNP-VP further reduced macrophage accumulation despite no significant change in plaque size. Across both settings, MoNP-VP increased fibroblast-like cell populations and collagen deposition accompanied by activation of TGFβ signaling, features consistent with a more stable plaque. Importantly, repeated MoNP-VP administration was well tolerated and elicited no overt systemic toxicity.

**Conclusion:** These findings bridge mechanistic insight and therapeutic efficacy in pre-established plaques by demonstrating that MoNP-VP remodels the inflammatory plaque microenvironment and promotes lesion stabilization, supporting its translational potential as a precision nanotherapy for atherosclerosis.

## INTRODUCTION

Atherosclerosis, arising from the progressive buildup of lipid-laden, inflammatory plaques within the arterial wall, narrows and hardens the vessel and more critically, carries the risk of sudden and catastrophic rupture, leading to thrombotic events such as myocardial infarction and ischemic stroke^1,2^. Despite advances of lipid-lowering and anti-platelet therapies, substantial cardiovascular mortality and morbidity remain, highlighting that plaque stabilization and regression remain unmet therapeutic objectives in atherosclerosis. Landmark trials such as CANTOS and COLCOT established that inflammatory pathway modulation can reduce cardiovascular events, validating inflammation as a therapeutic target in atherosclerosis^3,4^. However, whether systemic immunomodulation produces meaningful plaque stabilization or regression remains incompletely resolved, and safety concerns may limit the sustained, high-intensity anti-inflammatory activity needed to remodel established lesions. Achieving effective atherosclerosis management may therefore require complementing systemic risk-factor control with therapeutic agents that act directly on established lesions to suppress local inflammation and promote stabilizing vascular remodeling. However, lesion-targeted therapeutic strategies remain underdeveloped, constrained by both a limited understanding of actionable therapeutic targets that can directly modulate plaque development and a lack of effective delivery vehicles capable of selectively concentrating therapeutic agents within the lesion microenvironment.

Plaque development is driven by complex pathological interactions among vascular and immune cells within the vessel wall. In response to atherogenic stimuli, endothelial cells (ECs) become activated and dysfunctional, triggering infiltration of monocytes that differentiate into proinflammatory macrophages (Mφs) and form foam cells. In parallel, vascular smooth muscle cells (VSMCs) undergo phenotypic switching away from a contractile state, contributing to intimal thickening and lesion progression^5,6^. Despite extensive studies of the molecular mechanisms underlying atherosclerosis, therapeutic targets capable of coordinately modulating these multicellular pathological processes remain poorly defined. Emerging evidence, including ours, has identified yes-associated protein (YAP) and transcriptional co-activator with PDZ-binding motif (TAZ) as key regulators shared across multiple atherogenic processes in the vessel wall^7–13^. Specifically, endothelial YAP/TAZ hyperactivation promotes a pro-inflammatory phenotype that drives monocyte recruitment and early lesion development^7–9^. Beyond ECs, YAP/TAZ signaling promotes Mφ inflammatory polarization and chemokine secretion^10,11^, while in VSMCs, YAP/TAZ activation drives synthetic phenotype switching and medial remodeling, collectively accelerating plaque progression and instability^12,13^. These findings position YAP/TAZ as a shared regulatory axis across multiple disease-driving cell populations and suggest that lesion-targeted inhibition of YAP/TAZ-mediated transcription may coordinately suppress multicellular plaque pathology. However, translating this therapeutic potential requires selective inhibition of YAP/TAZ activity within diseased vascular lesions while limiting systemic exposure.

To address this challenge, we recently developed monocyte membrane-cloaked nanoparticles (MoNP) that exploit the natural inflammatory tropism of monocytes to home to inflamed vascular endothelium and accumulate within atherosclerotic lesions^14,15^. By encapsulating verteporfin (VP), a pharmacological inhibitor of YAP/TAZ, we generated MoNP-VP and demonstrated that it suppressed YAP/TAZ-mediated transcription in ECs and attenuated cytokine-induced pro-inflammatory activation. Importantly, MoNP-mediated delivery of VP, but not free VP, significantly reduced plaque formation in the partial carotid ligation (PL) model in apolipoprotein E-deficient (ApoE^-/-^) mice. These findings established the lesion-targeted, pathway-modulating activity of MoNP-VP, as well as its efficacy in suppressing endothelial inflammation and early plaque formation. However, the pharmacodynamic effects of MoNP-VP within the broader plaque microenvironment, particularly in Mφs and VSMCs where YAP/TAZ activation also contributes to disease progression, remain unknown. Moreover, reduced plaque formation represents an early proof of concept; whether MoNP-VP can halt progression and promote stabilizing remodeling of existing plaques remains a more clinically relevant and unresolved question.

In the present study, we combined single-cell RNA sequencing (scRNA-seq) with the ApoE^-/-^PL model to define the multicellular pharmacodynamics of MoNP-VP within the plaque microenvironment, extending beyond its established role in suppressing endothelial inflammation. This analysis identified MoNP-VP-associated shifts in lesion cellular composition at single-cell resolution. We then extended therapeutic evaluation to the adeno-associated virus (AAV)-PCSK9 model of hyperlipidemia-driven atherosclerosis, using a long-term diet-switch design in which mice either continued a high-fat diet (HFD) or switched to chow to assess the effects of MoNP-VP on established plaque progression under persistent versus reduced lipid burden. These studies demonstrate that MoNP-VP suppresses Mφ-driven inflammation, promotes VSMC/fibroblast-associated remodeling and collagen deposition, and activates transforming growth factor β (TGFβ) signaling. Collectively, our findings identify MoNP-VP as a lesion-directed nanotherapeutic that limits plaque progression by suppressing local inflammation while promoting stabilizing vascular remodeling.

## MATERIALS AND METHODS

### Nanoparticle Preparation

VP loaded nanoparticles (NP-VP) and nanoparticles (NP) were formulated using a single emulsion method as previously published^14^. 10 mg of PLGA (Resomer® RG 503H, Sigma Aldrich) with or without 2 mg of VP (Tocris Bioscience) was dissolved in dichloromethane (Sigma Aldrich) and subsequently introduced dropwise and emulsified in a 2% polyvinyl-alcohol (PVA, Acros Organics) solution, followed by solvent evaporation in 0.5% PVA. The formulated NP-VP and NP were collected by centrifugation. Monocyte membranes, isolated from mouse bone marrow-derived monocytes using a membrane isolation kit (Invent Biotechnologies), were mixed with nanoparticles at a 1:10 weight ratio and sonicated to form MoNP-VP and MoNP, respectively. Nanoparticle properties were characterized by dynamic light scattering (Zetasizer, Malvern) and transmission electron microscopy (Talos L120C, Thermo FisherScientific) at ASU Eyring Materials Center. For encapsulation efficiency and loading capacity measurements, NP-VP were dissolved in DMSO, and fluorescence intensity was measured using a plate reader (BioTek) following a previously reported method.

### Mouse models of Atherosclerosis

All animal experiments were approved by the Arizona State University Institutional Animal Care and Use Committee (IACUC; protocol nos. 23-1960R and 26-2173R). For the scRNA-seq analysis, ApoE^-/-^mice fed a HFD underwent PL of the left carotid artery (LCA) under isoflurane anesthesia (3-4% for induction and 1-3% for maintenance), which local disturbed flow patterns to induce endothelial activation and acute atherosclerosis^14,16^. Following surgery, mice were intravenously administered either MoNP-VP (VP dose: 2 mg/kg) or MoNP every 72 hours for a total of 3 injections. Ten days post-surgery, mice were euthanized by CO_2_ inhalation followed by cervical dislocation. Mice were then perfused with PBS to remove blood, and the collected vessels were allocated for either Oil Red O (ORO) staining (Sigma Aldrich) or scRNA-seq.

The effect of MoNP-VP was also evaluated in mice with pre-existing plaques. Briefly, C57BL/6J mice received a single intravenous injection of AAV-PCSK9 (5 × 10^11^ GC; Vector Biolabs) and were subsequently fed a HFD for 16 weeks to induce aortic atherosclerosis. After 16 weeks, a subset of mice was euthanized as the baseline group, while the remaining mice were randomly divided into two cohorts: one maintained on HFD (HFD-HFD) and the other switched to a chow diet (HFD-Chow) for an additional 6 weeks. During this period, mice in both cohorts received either MoNP-VP (VP dose: 2 mg/kg) or MoNP every 72 hours for a total of 12 injections. At the study endpoint, mice were euthanized by CO_2_ inhalation followed by cervical dislocation, and aortas, carotid arteries, major organs, and serum samples were collected for further analyses. Total cholesterol and metabolic panel analyses were performed by Vetek Lab (Scottsdale, Arizona). ORO staining was used to evaluate lesion burden in the whole arterial tree and aortic root cryosections, and lesion areas were quantified using ImageJ. For immunofluorescence staining, aortic root cryosections were incubated with antibodies against CD68 (#137001, BioLegend, 1:200), TREM2 (ab245227, Abcam, 1:200), CTSB (#31718S, Cell Signaling Technology, 1:200), FSP1 (16105-1-AP, Proteintech, 1:200), αSMA (#48938S, Cell Signaling Technology, 1:200), and pSMAD2/3 (PA5-110155, Thermo Fisher Scientific,1:200). Collagen content was assessed using Masson’s trichrome staining (#25088, Polysciences). Hematoxylin and eosin (H&E) staining was performed on liver sections to assess toxicity. Images were acquired using a BioTek Lionheart microscope and an Olympus VS200 slide scanner. Quantification was performed using ImageJ or QuPath.

### scRNA-seq analysis

LCAs collected from PL mice treated with MoNP-VP or MoNP were enzymatically digested using a cocktail containing hyaluronidase (0.8 mg/mL, Sigma-Aldrich), Liberase (2 U/mL, Sigma-Aldrich), and DNase I (60 U/mL, Sigma-Aldrich) to obtain single-cell suspensions. Cell number and viability were assessed using a Countess 3FL cell counter. Single cell libraries were prepared using the Chromium Single Cell 3’ Gene Expression v3.1 kit (10x Genomics) targeting approximately 5,000 cells per sample according to the manufacturer’s instructions. Sequencing was performed on an Illumina NovaSeq platform with an average depth of approximately 50,000 reads per cell.

Raw FASTQ files were processed using Cell Ranger v7.0.1 (10x Genomics) for read demultiplexing, alignment to the mouse reference genome (mm10-2020-A), and generation of gene-cell count matrices based on unique molecular identifier quantification. Count matrices from all samples were subsequently imported into Seurat v5 for QC and downstream analysis following the scSignalMap pipeline^17^. Cells with fewer than 5,000 counts per gene or greater than 8% mitochondrial gene expression were excluded to remove low-quality or apoptotic cells, and genes detected in only a limited number of cells were filtered out. Additionally, we excluded cells with more than 190,000 or 150,000 counts per cell to remove multiplets in the MoNP and MoNP-VP samples, respectively. A total of 2,064 cells from four LCAs in the MoNP-VP group and 2,850 cells from three LCAs in the MoNP group were included in downstream analyses. Data were normalized and scaled using SCTransform. The two samples were integrated using the IntegrateData Seurat method with SCT-normalized data. Marker genes were discovered using the Wilcoxon rank-sum test with the PrepSCTFindMarkers and FindAllMarkers functions in Seurat. Cell types were determined based on overlap with canonical marker gene expression. The integrated dataset was visualized using UMAP. The scSignalMap package (https://github.com/plaisier-lab/scSignalMap) was used to discover ligand-receptor pairs between cell types and pathways being differentially expressed downstream of the receptor in receiving cells^17^. The scSignalMap results were imported into a Neo4j graph database to enable interactive querying and visualization of treatment-specific changes in intercellular communication between the MoNP and MoNP-VP groups. The raw and processed single-cell RNA sequencing data have been deposited in the GEO under accession number GSE337236.

### Cell culture and *in vitro* assays

All human cells used in this study were obtained from commercial sources. Human aortic ECs (HAECs; #304-05a, Cell Applications) and human coronary artery smooth muscle cells (HCASMCs; #3514-05a, Cell Applications) were cultured in endothelial and smooth muscle cell growth medium, respectively. THP-1 monocytes (TIB-202^TM^, ATCC) were cultured in RPMI 1640 medium supplemented with 10% fetal bovine serum (FBS; #89510-186, Avantor). All cells were maintained in a humidified incubator at 37°C with 5% CO_2_.

For endothelial inflammation studies, ECs were pre-treated with MoNP-VP (VP dose: 1 μM) or MoNP for 3 hours, followed by TNFα stimulation (1 ng/mL) for an additional 3 hours. Cells were subsequently collected for real-time PCR analysis or subjected to monocyte adhesion assays to evaluate anti-inflammatory effects. For real-time PCR, total RNA was isolated using the Direct-zol RNA Kit (Zymo Research) and reverse transcribed using oligo(dT) primers and M-MLV reverse transcriptase (Promega). Quantitative real-time PCR was performed using a CFX Duet Real-Time PCR System (Bio-Rad). Relative gene expression levels were calculated using the 2^−ΔΔCT^ method. Primer sequences are listed in Supplementary Information. For monocyte adhesion assays, THP-1 monocytes were labeled using CellBrite® Cytoplasmic Membrane Dye (Biotium) and added to TNFα-treated ECs pretreated with MoNP-VP or MoNP at a concentration of 5 × 10^5^ cells/mL. After 30 minutes of incubation, unbound THP-1 monocytes were removed, and adherent monocytes were imaged using fluorescence microscopy and quantified using ImageJ.

For foam cell formation assays, THP-1 monocytes were differentiated into THP-1-derived Mφ using phorbol 12-myristate 13-acetate (PMA; 100 ng/ml) overnight and subsequently treated with MoNP-VP (VP dose: 1 μM) or MoNP for 3 hours. Oxidized low-density lipoprotein (oxLDL) (15 μg/ml) was then added to induce foam cell formation for 24 hours. Cells were fixed and stained with ORO to assess lipid accumulation. For EdU incorporation assays, HCASMCs were first treated either MoNP-VP (VP dose: 1 μM) or MoNP for 3 hours and then stimulated with PDGF-BB (20 ng/ml) for 48 hours. EdU reagent was added 2 hours prior to cell fixation. EdU staining was performed according to the manufacturer’s protocol.

### Statistical analysis

All experiments were performed with at least three biological replicates, and data are presented as mean ± standard deviation (SD). Statistical analyses were conducted using GraphPad Prism 10.5. Comparisons between two groups were performed using Student’s t-test, while comparisons among multiple groups were analyzed using one-way ANOVA followed by Tukey’s multiple comparisons test.

## RESULTS

### MoNP-VP suppresses pro-atherogenic vascular remodeling at single-cell resolution

MoNP-VP was prepared as previously described^14^. Consistent with our prior report, MoNP-VP exhibited the expected physicochemical properties and VP encapsulation, and MoNPs preferentially accumulated in atherosclerotic arteries (**Supplementary Figure 1**). To evaluate the multicellular effects of MoNP-VP on the atherosclerotic microenvironment, we performed PL of the LCA in ApoE^-/-^mice fed a HFD. MoNP-VP (2 mg/kg of VP) or empty MoNP vehicle control was administered intravenously via retro-orbital injection every 72 hours for a total of three injections. Mice were euthanized 10 days post-PL to assess early lesion formation, and carotid arteries were harvested for histological analysis and scRNA-seq profiling (**Figure 1A**). Whole-mount tissue imaging and ORO staining of LCA sections confirmed plaque formation in MoNP-treated mice, whereas no detectable luminal lesions were observed in MoNP-VP-treated mice (**Figure 1B & 1C**), further supporting our previous findings^14^. To profile cellular populations within MoNP-VP- and MoNP-treated arterial walls, LCAs from a separate cohort were pooled within each treatment group, enzymatically dissociated into single-cell suspensions, and processed for droplet-based scRNA-seq. After quality control filtering and doublet removal, 2,064 cells from MoNP-VP-treated and 2,850 cells from MoNP-treated LCAs were retained for downstream analysis (**Supplementary Figure 2**). Cells from the two treatments were integrated and *de novo* clustering identified nine transcriptionally distinct clusters present in both groups (**Figure 1D**). Clusters were annotated based on established marker genes, identifying VSMC/fibroblasts (VSMC/FB) and endothelial cells (EC), as well as immune cell populations, including four monocyte/Mφ clusters (Mo1-2 and Mφ1-2), granulocytes (GR), dendritic cells (DC), and T cells (T)^18–25^ (**Figure 1E**). Notably, a transcriptionally distinct VSMC cluster was not readily identifiable; instead, the predominant stromal population exhibited a fibroblast-enriched transcription profile with relatively low expression of canonical contractile VSMC markers. We therefore annotated these cells as a unified VSMC/FB cluster based on their overlapping transcriptional signature. Among the four monocyte/Mφ populations, Mo1 expressed classical monocyte markers *Ly6c2* and *Ccr2*, while Mo2 expressed genes more closely associated with non-classical monocytes, including *Cxcr4* and *Fcgr4*, with low *S100a8* and *S100a9* and absent *Ly6c2* and *Ccr2* expression. Mφ1 and Mφ2 showed the most defined functional profiles, consistent with inflammatory and foam cell-associated Mφ states characterized by *Tnf* and *Nlrp3*, and *Trem2* and *Ctsb* expression, respectively^23,25^. Violin plots of additional marker genes are provided in **Supplementary Figure 3** to further illustrate the transcriptional characteristics of the annotated cell populations.

**Figure 1.**
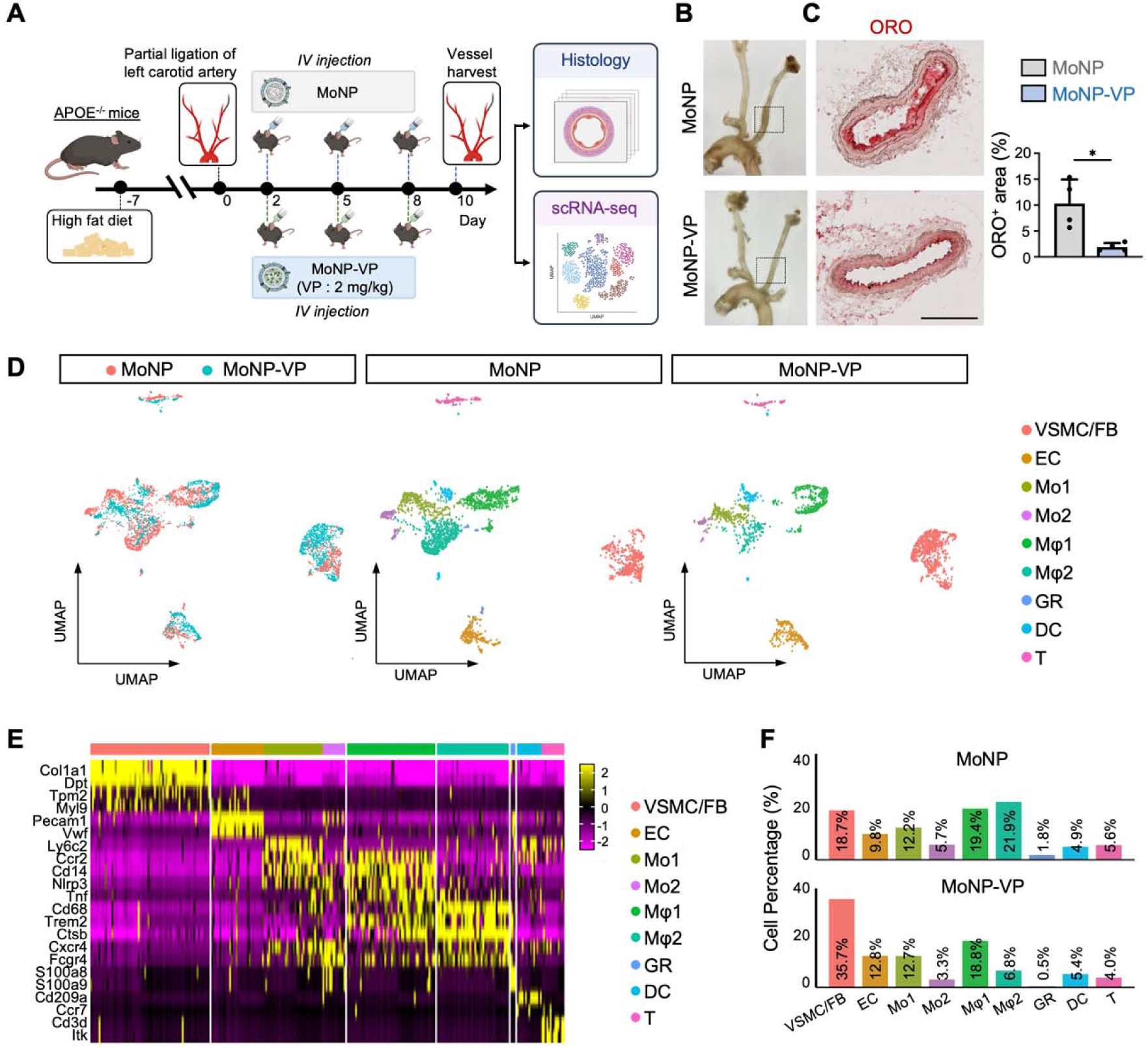
scRNA-seq profiling of partially ligated left carotid atherosclerotic arteries following MoNP-VP treatment. (A) Schematic diagram of the animal experiment. ApoE^-/-^mice were fed a HFD and underwent partial ligation surgery, followed by treatment with MoNP-VP or MoNP. Vessels were collected for histological assessment and scRNA-seq profiling. (B-C) Representative vessel images (B) and ORO-stained cross sections of the LCA (C) showing lesion formation in the MoNP-VP and MoNP groups, along with quantification results. Scale bar = 200 μm. n = 4. (D) UMAP visualization showing nine identified cell clusters in both the MoNP-VP and MoNP groups. (E) Heatmap showing representative marker genes for each cluster. (F) Bar graphs showing the cell counts and cell percentages of each cluster in the MoNP-VP and MoNP groups.

Comparison of cluster proportions between MoNP-VP- and MoNP-treated LCAs revealed marked shifts in lesion cellular composition (**Figure 1F**). The most prominent treatment-associated changes were observed in the VSMC/FB and Mφ2 clusters. The VSMC/FB population increased from 18.7% in MoNP-treated arteries to 35.7% following MoNP-VP treatment, whereas the Mφ2 foam cell-associated Mφ population was markedly reduced from 21.9% to 6.8%. Other cell populations showed more modest differences between MoNP-VP and MoNP groups, including EC (12.8% vs. 9.8%), Mo1 (12.7% vs. 12.2%), Mo2 (3.3% vs. 5.7%), Mφ1 (18.8% vs. 19.4%), GR (0.5% vs. 1.8%), DC (5.4% vs 4.9%), and T (4.0% vs 5.6%), respectively. Together, these findings demonstrate that MoNP-VP remodels the atherosclerotic vessel wall beyond its previously established effects on endothelial inflammation, primarily by reducing foamy Mφ accumulation and expanding VSMC/FB-associated stromal populations.

### MoNP-VP limits progression of pre-existing aortic lesions

Having identified MoNP-VP-associated remodeling of lesion cellular composition in the ApoE^-/-^PL model, we next evaluated whether these effects translated into therapeutic benefit in established plaques. Using an AAV-PCSK9-induced atherosclerosis model^26^, we assessed whether MoNP-VP treatment could limit progression of pre-existing lesions under persistent hypercholesterolemia or after dietary cholesterol lowering. C57BL/6J mice received 5×10¹¹ viral particles of AAV-PCSK9 and were fed a HFD for 16 weeks to induce aortic plaque formation. Saline-injected mice maintained on HFD served as a non-atherosclerotic control, confirming that lesion formation was driven by AAV-PCSK9-induced hypercholesterolemia (**Supplementary Figure 4**). After 16 weeks of HFD, a subset of AAV-PCSK9-treated mice was harvested to establish baseline pre-existing plaque burden, while the remaining mice were randomized to either continue HFD (HFD-HFD) or switch to chow diet (HFD-Chow) for an additional 6 weeks. During this period, mice received MoNP-VP (2 mg/kg) or MoNP twice weekly for a total of 12 injections (**Figure 2A**). At the end of the 22-week study period, serum and arterial tissues were collected for lipid measurement and lesion analysis. Total cholesterol and triglyceride levels remained elevated in HFD-HFD mice compared with the 16-week baseline group, whereas HFD-Chow mice showed marked reduction in both lipid measures after diet switch (**Figure 2B**), confirming persistent hypercholesterolemia under continued HFD and effective cholesterol lowering after dietary intervention. Lesion burden was assessed by ORO staining of whole-mount arterial tissues spanning from the iliac bifurcation through the abdominal and thoracic aorta, aortic arch, and carotid arteries (**Figure 2C & 2D**), as well as cross-sections of the aortic root (**Figure 2E & 2F**). At the 16-week baseline, established lesions were present in both whole-mount aortas and aortic root sections. In HFD-HFD mice, MoNP-treated controls showed significant lesion progression beyond baseline, whereas MoNP-VP significantly attenuated lesion growth across both anatomical readouts. In HFD-Chow mice, lesion burden remained comparable to baseline in both MoNP- and MoNP-VP-treated groups, indicating that dietary cholesterol lowering was sufficient to limit further plaque growth; in this cholesterol-lowering setting, MoNP-VP did not further reduce bulk lesion area compared with MoNP. These results demonstrate that MoNP-VP effectively attenuates progression of established lesions under continued HFD, while dietary cholesterol lowering limits further plaque progression in the HFD-Chow setting.

**Figure 2.**
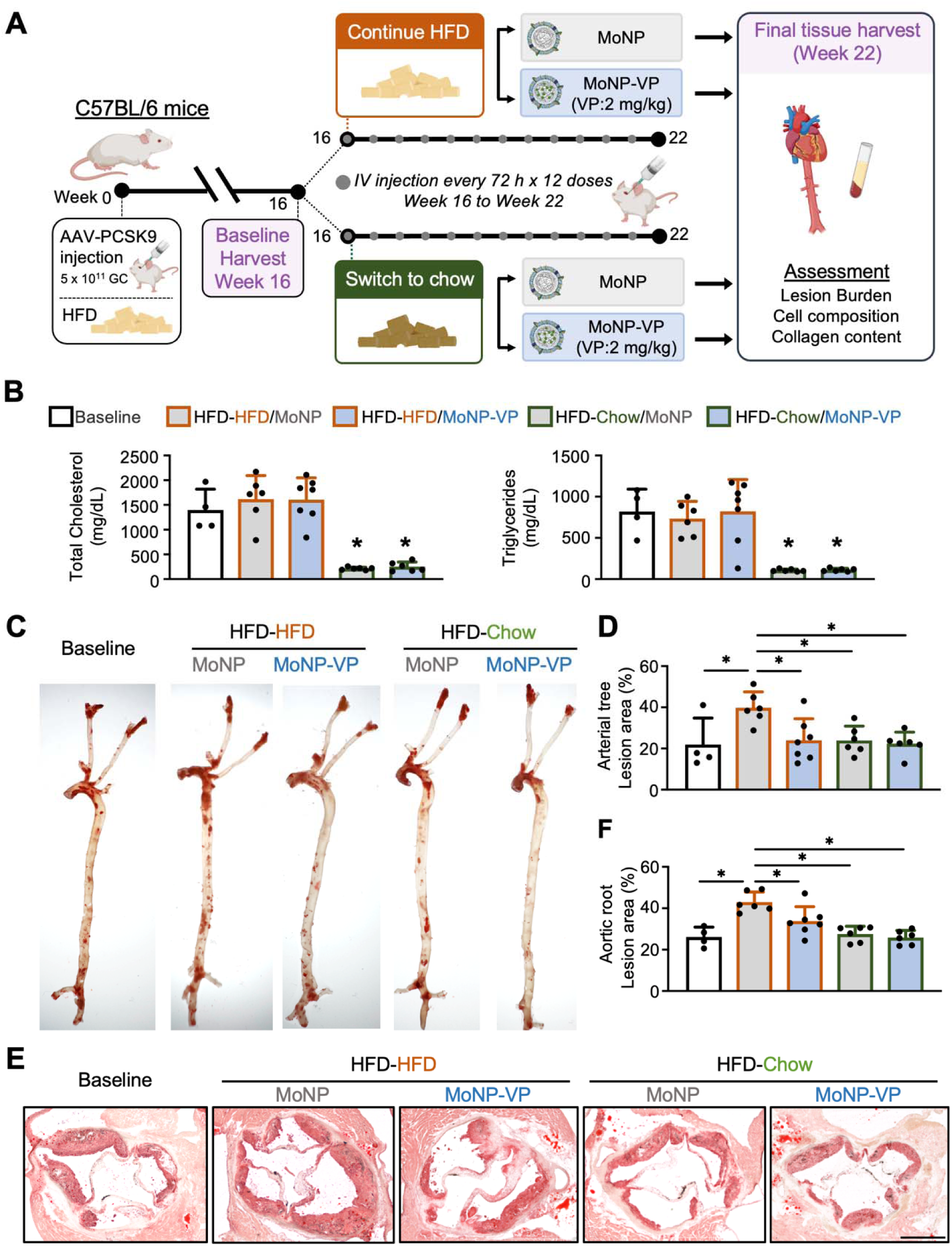
MoNP-VP attenuated plaque progression in established plaques. (A) Schematic diagram of the animal experiment. C57BL/6 mice received AAV-PCSK9 and were fed a HFD for 16 weeks to induce atherosclerosis. A subset of mice was euthanized at week 16 as the baseline group. The remaining mice were then randomized to either HFD-HFD or HFD-Chow, while receiving MoNP-VP or MoNP twice weekly for a total of 12 injections. At the endpoint, mice were euthanized, and organs and serum were collected for further assessment. (B) Bar graph showing total cholesterol levels and triglycerides each group. *p < 0.05 compared with the baseline and HFD-HFD groups. (C–F) Representative ORO staining images and quantification of the carotid arteries, whole aortas (C,D) and aortic root sections (E,F). (E) Scale bar = 500 μm. Baseline n = 4, HFD-HFD-MoNP-VP n = 7, and all other groups n = 6. *p < 0.05.

### MoNP-VP restrains Mφ accumulation and foam-cell-associated remodeling

To determine whether the Mφ compositional changes identified by scRNA-seq were reflected at the tissue level, we performed immunostaining for Mφ-associated markers in aortic root sections from the AAV-PCSK9 diet-switch model. Guided by the observed reduction in the Mφ2 cluster, we used CD68 to assess overall Mφ burden and TREM2 and CTSB to evaluate foamy Mφ populations. In HFD-HFD mice, MoNP-VP significantly reduced CD68+ area compared with MoNP controls (**Figure 3A**). In HFD-Chow mice, dietary cholesterol lowering alone reduced CD68+ area relative to HFD-HFD MoNP controls, while MoNP-VP further decreased CD68+ area compared with MoNP treatment, indicating an additional effect of MoNP-VP on Mφ burden under cholesterol-lowering conditions. For foam cell-associated markers, MoNP-VP significantly reduced CTSB+ area and showed a trend toward reduced TREM2+ area under continued HFD conditions (**Figure 3B & 3C**). In HFD-Chow mice, CTSB and TREM2 levels were comparable between MoNP-VP and MoNP groups, suggesting that the effect of MoNP-VP on these foamy Mφ markers was less apparent after dietary cholesterol lowering. These findings support the scRNA-seq results and indicate that MoNP-VP reduces overall Mφ burden, with its strongest effects on foamy Mφ markers observed under persistent hypercholesterolemic stress.

**Figure 3.**
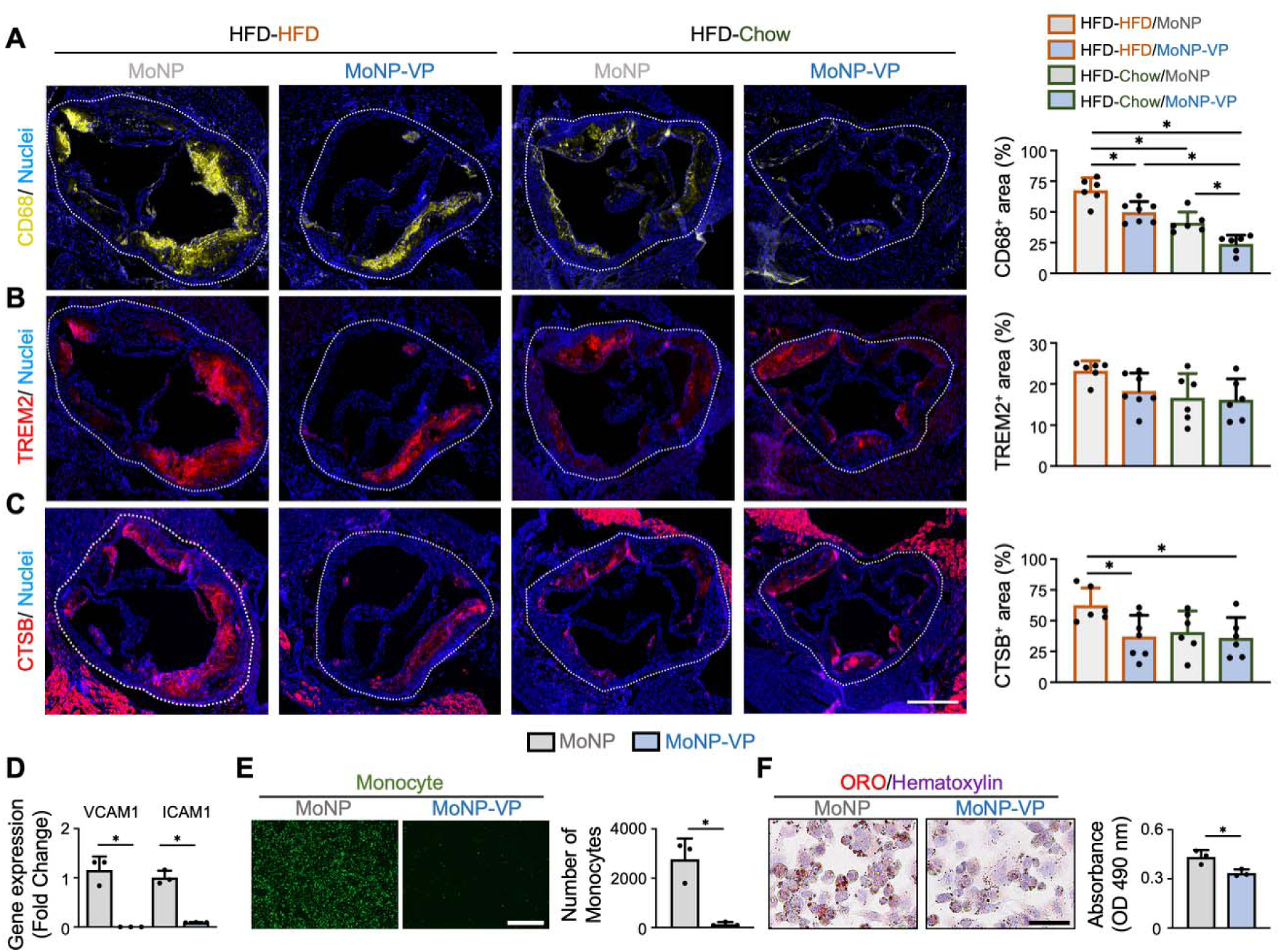
MoNP-VP suppresses inflammatory plaque progression. (A–C) Representative immunostaining images and quantification of CD68+ (A), TREM2+ (B), and CTSB+ (C) areas in atherosclerotic plaques. Scale bar = 400 μm. Baseline n = 4, HFD-HFD-MoNP-VP n = 7, and all other groups n = 6. *p < 0.05. (D) Gene expression levels of VCAM1 and ICAM1 in TNFα-stimulated ECs with MoNP-VP or MoNP pretreatment. (E) Representative images and quantification of monocyte adhesion to TNFα-stimulated ECs with MoNP-VP or MoNP pretreatment. Scale bar = 500 μm. (F) ORO staining and quantification of foam cell formation in oxLDL-stimulated THP1-derived Mφ with MoNP-VP or MoNP pretreatment. Scale bar = 50 μm. D–F, n = 3. *p < 0.05.

To further define potential mechanisms, we performed *in vitro* studies using HAECs and THP-1 cells. Consistently, MoNP-VP reduced endothelial adhesion molecule expression and decreased monocyte recruitment under inflammatory stimulation (**Figure 3D & 3E**). In THP-1-derived Mφs, MoNP-VP reduced oxLDL uptake compared with MoNP control, suggesting a direct effect on foam cell formation (**Figure 3F**). Together, these tissue-level and *in vitro* findings indicate that MoNP-VP suppresses Mφ-associated plaque burden through complementary mechanisms, including reduced endothelial-mediated monocyte recruitment and direct attenuation of Mφ lipid uptake.

### MoNP-VP induces fibroblast-like stromal enrichment and collagen-rich plaque stabilization

We next assessed stromal remodeling following MoNP-VP treatment in aortic root sections from the AAV-PCSK9 diet-switch model by staining for the fibroblast marker FSP1 and the contractile VSMC marker αSMA. MoNP-VP significantly increased FSP1+ cell content compared with MoNP controls in HFD-HFD mice, with a similar increase observed in HFD-Chow mice (**Figure 4A**), consistent with expansion of the VSMC/FB cluster identified by scRNA-seq. In contrast, αSMA+ cell content was not significantly different between treatment groups under either dietary condition, suggesting that MoNP-VP promotes fibroblast-like stromal enrichment rather than restoring the contractile VSMC population, which is diminished in atherosclerotic arteries. To assess whether the increased FSP1+ stromal content could be explained by enhanced VSMC proliferation, we examined EdU incorporation in PDGF-BB-stimulated human coronary artery VSMCs *in vitro*. MoNP-VP reduced the number of EdU-incorporated VSMCs compared with MoNP controls (**Figure 4B**), suggesting that the FSP1+ enrichment observed *in vivo* more likely reflects a fibroblast-like phenotypic shift rather than mitogenic VSMC expansion.

**Figure 4.**
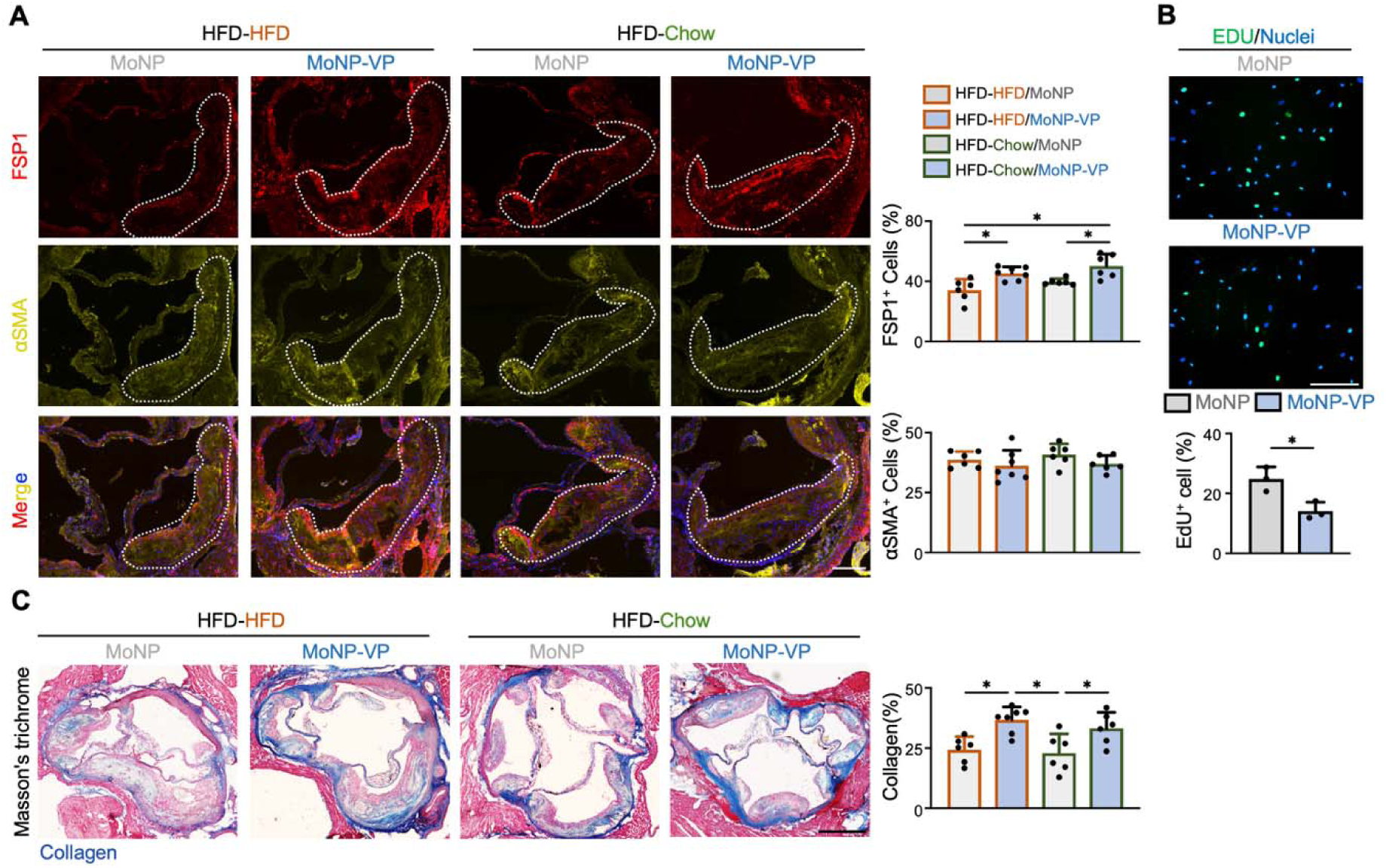
Changes in plaque cellular composition and collagen content following MoNP-VP treatment. (A) Representative immunostaining images and quantification of FSP1+ and αSMA+ content in the aortic root following MoNP-VP treatment under both conditions. Scale bar = 200 μm. (B) EdU assay and quantification of PDGFBB-stimulated VSMC proliferation following MoNP-VP or MoNP treatment. n = 3. *p < 0.05. Scale bar = 200 μm. (C) Representative Masson’s trichrome staining images and quantification of collagen content in the aortic root following MoNP-VP treatment under both conditions. Scale bar = 400 μm. (A,C) Baseline n = 4, HFD-HFD-MoNP-VP n = 7, and all other groups n = 6. *p < 0.05. Data are presented as mean ± SD.

Because collagen deposition is a key determinant of plaque mechanical stability and is produced by fibrogenic stromal cells including fibroblast-like VSMCs and resident fibroblasts^27^, we next assessed collagen content by Masson’s trichrome staining. MoNP-VP significantly enhanced collagen deposition compared with MoNP controls under both dietary conditions (**Figure 4C**). These findings demonstrate that MoNP-VP promotes fibroblast-like stromal enrichment and collagen-rich matrix remodeling in established atherosclerotic lesions, collectively supporting a shift toward a more mechanically stable plaque composition.

### MoNP-VP-induced plaque remodeling is associated with enhanced TGFβ/SMAD signaling

To investigate intercellular signaling mechanisms underlying MoNP-VP-associated plaque remodeling, we identified ligand-receptor interactions using the scRNA-seq data by pairing ligands expressed by sending cell populations with receptors upregulated in receiving cell populations to infer downstream signaling pathways (**Figure 5A**)^17^. The resulting interactions across multiple cell types within the plaque microenvironment were visualized as a network, revealing broad MoNP-VP-associated changes in intercellular communication (**Figure 5B**). A more focused look at the most significantly differentially upregulated receptors (log_2_FC ≥ 1) demonstrated that the TGFβ-TGFβR axis was one of the most prominent signaling pathways (**Figure 5C**). Downstream pathway analysis of the ligand-receptor network further revealed enrichment of TGFβ-associated biological processes, including vascular development, collagen fibril organization, extracellular matrix (ECM) organization, and TGFβ receptor signaling (**Figure 5D**), consistent with the established role of TGFβ signaling in promoting plaque-stabilizing remodeling^28–30^. Based on these results, we examined phosphorylation of SMAD2/3 (pSMAD2/3) as a canonical downstream readout of TGFβ pathway activation. In the pre-existing plaque model, pSMAD2/3+ staining was significantly elevated in MoNP-VP-treated aortic roots compared with MoNP controls in the HFD-HFD group (**Figure 5E**), supporting TGFβ pathway activation in response to MoNP-VP. In the HFD-Chow group, however, no significant difference in pSMAD2/3 staining was observed between MoNP-VP and MoNP groups, suggesting MoNP-VP-associated TGFβ signaling may be less pronounced under cholesterol-lowering conditions. Together, these findings indicate that MoNP-VP enhances TGFβ signaling most prominently under conditions of active plaque progression, supporting TGFβ-associated intercellular signaling as a potential mechanism contributing to MoNP-VP-mediated plaque remodeling.

**Figure 5.**
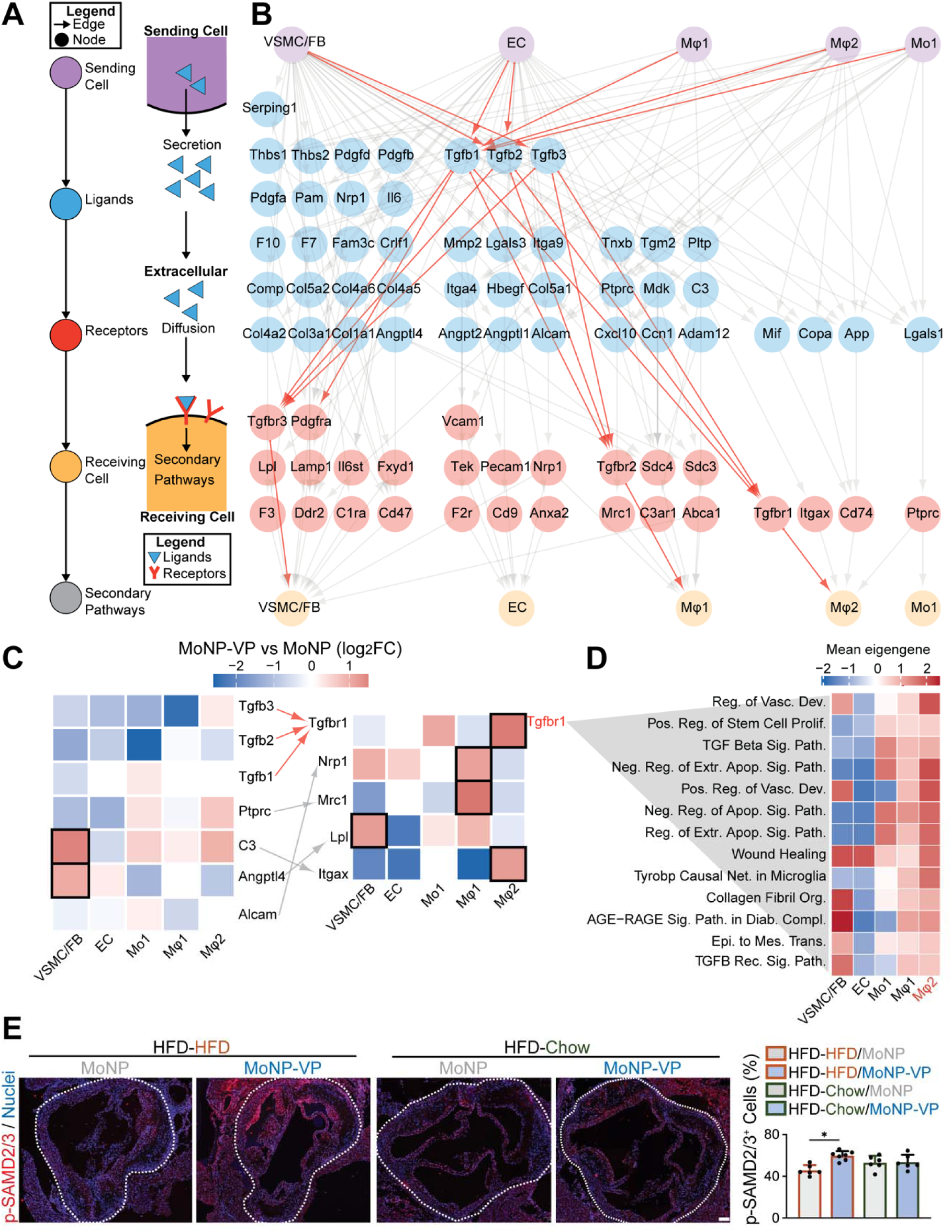
MoNP-VP remodels the plaque microenvironment through TGFβ pathway activation. (A) Schematic of ligand-receptor-mediated intercellular signaling. (B) Network generated by scSignalMap from scRNA-seq data showing ligands interacting with receptors upregulated by MoNP-VP relative MoNP. Nodes represent biological entities, and directed edges indicate interactions. TGFβ pathway related interactions are highlighted in red. (C) Receptors from (B) with log_2_FC ≥ 1 are shown with ligands (left) and receptors (right) for each cluster from the subset. Red indicates expression level upregulation and blue indicates downregulation. Arrows indicate directed relationships for ligand-receptor pairs, TGFβ pathway is highlighted in red. (D) Secondary pathway analysis of TGFβR1 for pathways chosen with highest gene overlap percent (5-11%). (E) Representative immunostaining and quantification of p-SMAD2/3⁺ cells in the aortic root following MoNP-VP or MoNP treatment. Scale bar = 100 μm.Baseline n = 4, HFD-HFD-MoNP-VP n = 7, and all other groups n = 6. *p < 0.05. Data are presented as mean ± SD.

### Repeated MoNP-VP treatment shows no detectable systemic or hepatic toxicity

To assess tolerability of the repeated treatment regimen, body weight was monitored throughout the 6-week treatment period in both HFD-HFD and HFD-Chow mice, and serum biochemistry and liver histology were evaluated at the study endpoint. No substantial differences in body weight were observed between MoNP-VP- and MoNP-treated mice in either dietary group at endpoint (**Figure 6A**). Serum biochemical analysis, including markers of hepatic function (ALP, ALT, AST, albumin, total protein, and total bilirubin), renal function (BUN and creatinine), and metabolic status (glucose and electrolytes), revealed no significant differences among baseline, MoNP-treated, and MoNP-VP-treated mice in the HFD-HFD group, indicating no detectable systemic metabolic, hepatic, or renal toxicity following repeated treatment (**Figure 6B, Supplementary Table 1**). In HFD-Chow mice, several hepatic function marker levels were reduced after diet switching, likely reflecting improved metabolic status following cholesterol lowering. Given the expected hepatic clearance of PLGA-based MoNP-VP^14^, we further performed H&E staining of liver sections to assess potential hepatotoxicity. Histological analysis showed preserved hepatic architecture with no apparent pathological abnormalities or tissue damage in MoNP-VP-treated mice compared with MoNP controls in both HFD-HFD and HFD-Chow groups (**Figure 6C**). Together, these findings support the tolerability of repeated MoNP-VP administration at the tested dose and regimen, 2 mg/kg VP twice weekly for 6 weeks, with no detectable systemic toxicity under either continued hypercholesterolemia or cholesterol-lowering conditions.

**Figure 6.**
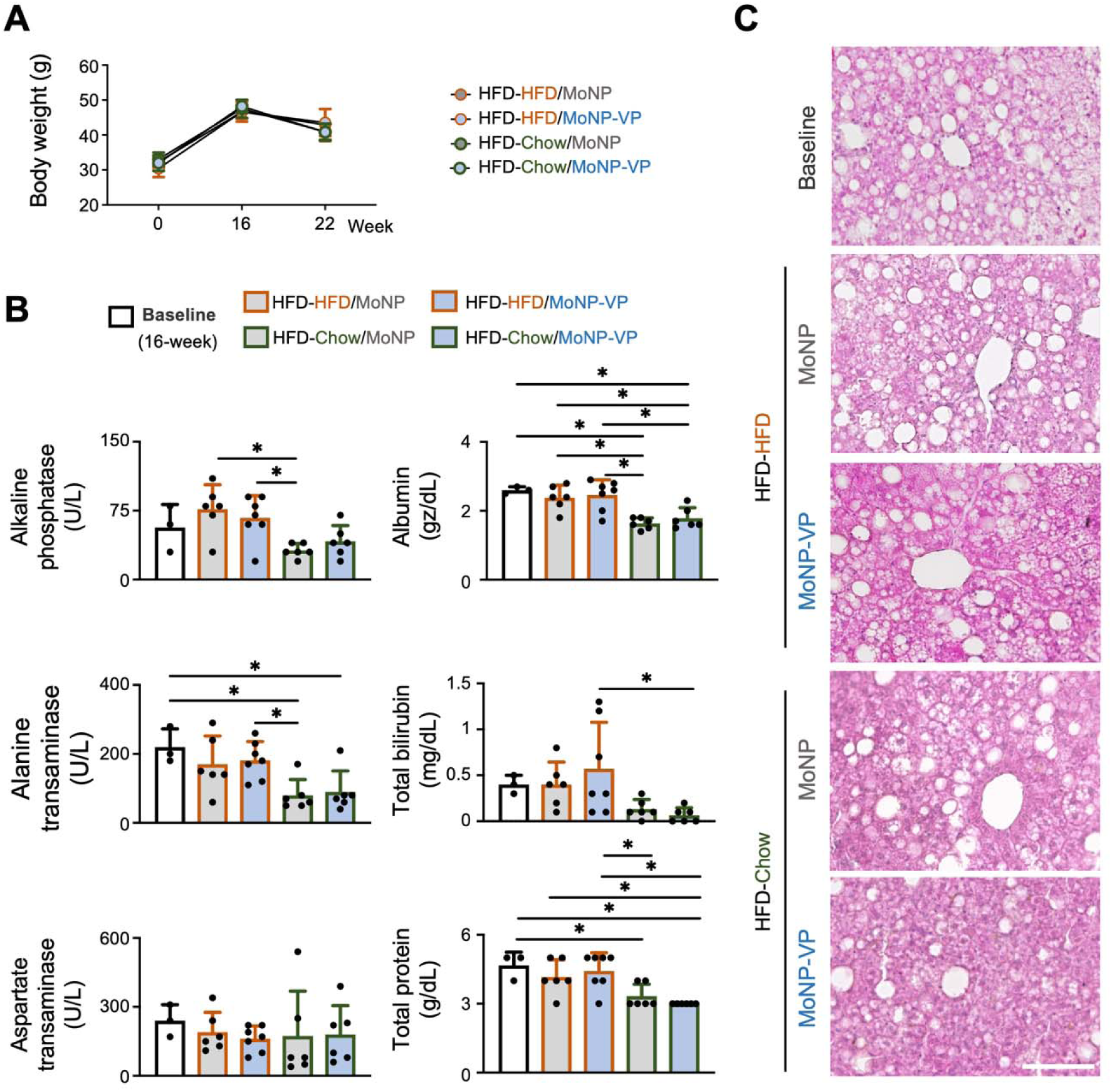
Biocompatibility analysis of MoNP-VP treatment. (A) Body weight measurements, (B) serum metabolic panel analysis, and (C) representative H&E staining images of liver sections following treatment. Scale bar = 100 μm. Baseline n = 3, HFD-HFD-MoNP-VP n = 7, and all other groups n = 6. *p < 0.05. Data are presented as mean ± SD.

## DISCUSSION

Atherosclerosis progression reflects complex interactions among dysfunctional ECs, dedifferentiated VSMCs, proinflammatory Mφs, and ECM components, a multicellular disease process that therapeutic strategies must coordinately address. Our previous work demonstrated that YAP/TAZ hyperactivation contributes to endothelial dysfunction and that MoNP-mediated delivery of VP reduces endothelial inflammation and attenuates plaque formation in the PL model. This present study extends the work by demonstrating that MoNP-VP effectively limits progression of pre-existing plaques through broader multicellular remodeling of the vessel wall. Specifically, MoNP-VP reduced foamy Mφ abundance, expanded fibroblast-like stromal populations, enhanced collagen deposition, and increased TGFβ/SMAD pathway activity within the lesion microenvironment. These effects were identified by scRNA-seq in partially ligated carotid arteries of ApoE^-/-^mice and corroborated by tissue-level analyses in a long-term AAV-PCSK9-driven atherosclerosis model. Together, these findings establish MoNP-VP as a lesion-directed nanotherapeutic that acts beyond its primary endothelial target to suppress Mφ-driven inflammation and promote collagen-rich matrix remodeling associated with plaque stabilization.

An important strength of this study is the evaluation of MoNP-VP efficacy across complementary models that capture distinct stages and lipid-burden contexts of disease. The ApoE^-/-^PL model combines disturbed flow with genetic hypercholesterolemia, producing spatially defined lesions within the LCA in less than 4 weeks and enabling rapid therapeutic screening and mechanistic assessment. The AAV-PCSK9 model, however, recapitulates key features of human familial hypercholesterolemia without requiring a germline knockout background^31–33^. Importantly, ApoE exerts immunomodulatory functions beyond lipoprotein clearance^34^; therefore, demonstrating therapeutic efficacy in a wild-type background driven by PCSK9 gain-of-function strengthens the translational relevance of MoNP-VP. The diet-switch design further enabled comparison between continued hypercholesterolemia and diet-induced lipid lowering, providing a clinically relevant framework to assess therapeutic benefit in established plaques across distinct progression states. Consistent with our previous findings in the ApoE^-/-^PL model^14^, the same MoNP-VP dose and treatment regimen that blocked plaque formation also limited progression of pre-existing plaques in AAV-PCSK9-treated mice maintained on HFD. However, MoNP-VP did not further reduce bulk plaque area in mice switched to chow. This may reflect, first, that dietary cholesterol lowering was sufficient to attenuate further plaque growth, thereby limiting the measurable additional effect of MoNP-VP on bulk lesion area. Second, reduced lipid burden may decrease endothelial activation, potentially diminishing MoNP-VP binding and accumulation at lesion sites. Nevertheless, these findings demonstrate that MoNP-VP efficacy is not limited to attenuating early atherogenesis but extends to limiting established plaque progression under persistent hypercholesterolemia, a scenario of considerable clinical relevance.

Plaque vulnerability depends not only on lesion size but also on inflammatory burden, vessel wall remodeling, and ECM composition, particularly collagen deposition^35,36^. In this study, MoNP-VP reduced Mφ accumulation while expanding VSMC/FB-associated stromal populations, supporting a shift toward a more stable plaque phenotype. This reduction in foam cell-associated Mφ population identified by scRNA-seq was further supported by decreased CD68, TREM2, and CTSB staining in the established plaque model, indicating that MoNP-VP attenuates inflammatory plaque burden beyond its effects on endothelial activation. Importantly, reduced CD68+ content was observed even in the HFD-Chow group, where MoNP-VP did not further reduce bulk lesion area. This suggests that lesion-targeted YAP/TAZ modulation may improve plaque composition even when dietary intervention is sufficient to limit further lesion growth. By comparison, effects on foamy Mφ markers TREM2 and CTSB were most evident under continued HFD conditions, suggesting that MoNP-VP has its strongest impact on foam-cell-associated Mφ states during active lipid-driven disease progression. More broadly, these findings further support that MoNP-VP as a potential complement to systemic cholesterol lowering by addressing residual inflammatory features of established plaques.

The reduction in overall Mφ burden may arise, at least in part, from decreased endothelial activation and diminished monocyte recruitment, consistent with our previous reports showing that YAP/TAZ inhibition by anti-sense oligos or MoNP-VP suppresses endothelial inflammation^7,14^. In addition, the preferential reduction of Mφ2 cluster suggests that VP released from MoNP may also act directly on lesional Mφs to reduce foam-cell formation or persistence. Supporting this possibility, our *in vitro* findings showed that MoNP-VP reduced oxLDL uptake in THP1-derived Mφs. Notably, in an unbiased screen of FDA-approved compounds, Hoeffner et al. identified VP as one of the most potent agents capable of reducing foam cell formation^37^, lending independent pharmacological support to our observations. This effect is mechanistically plausible because YAP/TAZ activity has been specifically implicated in driving M1 Mφ polarization and upregulating CD36-mediated oxLDL uptake through TEAD4^38^, two processes that together fuel foam-cell formation and plaque progression. Lesion-specific inhibition of YAP/TAZ by in the vessel wall by MoNP-VP may therefore provide a means of simultaneously suppressing Mφ inflammatory burden and reducing pathological lipid uptake. Together, these data suggest that MoNP-VP suppresses plaque development through complementary mechanisms: reduced monocyte recruitment following suppressed endothelial activation and potential direct modulation of foam cell formation programs through YAP/TAZ inhibition. Whether MoNP-VP directly alters Mφ pathways governing lipid uptake, cholesterol efflux, or efferocytosis warrants further investigation.

VSMCs are among the most abundant cell types in the arterial wall. During atherosclerosis development, VSMCs undergo extensive phenotypic modulation, characterized by downregulation of canonical contractile markers and transition into diverse synthetic states, including proliferative, Mφ-like, and fibroblast-like phenotypes^39,40^. Among these states, fibroblast-like VSMC-derived cells, termed fibromyocytes, are particularly relevant to plaque stability because they contribute to collagen production, fibrous cap formation, and ECM remodeling^41^. Consistent with this framework, MoNP-VP treatment was associated with expansion of the VSMC/FB cluster by scRNA-seq, corroborated histologically by increased FSP1+ cell content and enhanced collagen deposition within the vessel wall. However, αSMA+ population was unchanged, suggesting that MoNP-VP does not simply expand VSMCs but instead promotes enrichment of the fibromyocyte phenotype. Our *in vitro* findings support the interpretation that expansion of this population does not arise from VSMC proliferation but rather from fibroblast-like cells. Although the precise origin of these fibroblast-like cells cannot be determined from the current data, they may arise from phenotypic modulation of lesional VSMCs toward a fibromyocyte state, from expansion or migration of resident fibroblast populations, or from a combination of both. Nevertheless, MoNP-VP-induced enrichment of fibroblast-like cells and collagen-rich matrix within the lesion indicates a shift toward compositional features closely associated with plaque stabilization.

The reciprocal relationship between pro-inflammatory Mφ burden and VSMC/FB-associated remodeling is increasingly recognized as an important determinant of plaque vulnerability^41,42^. Accordingly, the interplay between these populations within the lesion microenvironment is of considerable mechanistic interest. Cell-cell communication analysis of our scRNA-seq data identified TGFβ-TGFBR pairs as a prominent intercellular axis associated with the expanded VSMC/FB cluster following MoNP-VP treatment. TGFβ signaling in atherosclerotic plaques is generally considered protective, in part by suppressing inflammation, promoting ECM deposition, and supporting fibrous cap integrity^42^. Consistent with this interpretation, immunostaining confirmed increased nuclear pSMAD2/3 staining, supporting activation of downstream TGFβ signaling after MoNP-VP treatment. Pro-inflammatory Mφs are known to suppress reparative plaque remodeling by secreting matrix metalloproteinases that degrade collagen and inflammatory cytokines, whereas reparative M2-like Mφs secret TGFβ to promote ECM remodeling^43,44^. Thus, a potential mechanistic link is that MoNP-VP reduces pro-inflammatory Mφ burden, creating a permissive microenvironment for TGFβ/SMAD signaling in adjacent VSMCs and fibroblast-like cells that promotes collagen-rich matrix remodeling and plaque stabilization. In addition, because VP is known to exert pharmacological effects beyond YAP/TAZ inhibition^45,46^, the observed remodeling may reflect combined effects of YAP/TAZ modulation, Mφ suppression, and broader VP-mediated changes in intercellular signaling. Together, these findings suggest that, beyond reducing endothelial activation and suppressing foam cell-associated programs, MoNP-VP reshapes the intercellular signaling landscape within the plaque, promoting TGFβ-driven atheroprotective remodeling in VSMCs and fibroblast-like cells.

Several limitations should be considered when interpreting these findings and in guiding future translational studies. First, all experiments were performed in male mice; whether MoNP-VP exerts comparable efficacy in female mice remains to be established, given well-documented sex differences in atherosclerotic plaque biology. Second, only a single VP dose formulation was evaluated. Although the tested dose of 2 mg/kg is substantially lower than doses reported in other disease models^47–49^, likely reflecting the enhanced delivery efficiency of the MoNP platform, systematic dose-response studies will be needed to define the therapeutic window and optimal dosing regimen of this approach. Third, although repeated MoNP-VP administration produced no measurable adverse effects in our toxicity analyses, suggesting tolerability even in the setting of metabolic burden, longer-term safety studies will be needed. In particular, future studies should evaluate whether sustained modulation of YAP/TAZ signaling has unintended effects on vascular remodeling in non-target organs and tissues.

## CONCLUSION

In summary, building on our previous work, this study further demonstrates that lesion-targeted, pathway-specific nanotherapy MoNP-VP not only attenuates early plaque formation but also limits the progression of pre-existing plaques in clinically relevant models of persistent dyslipidemia and dietary cholesterol lowering. Pharmacomechanistic studies integrating scRNA-seq analysis, histology, and complementary *in vitro* models reveal that MoNP-VP acts beyond suppression of endothelial inflammation; instead, it promotes multicellular remodeling of the atherosclerotic vessel wall, characterized by reduced total Mφ burden, attenuation of foam-cell-associated Mφ programs, expansion of fibroblast-like cell populations, and enhanced collagen deposition. Together, these changes indicate a shift toward a more stabilized plaque phenotype. These findings support MoNP-VP treatment as a potential complement to systemic cholesterol-lowering therapy by directly improving plaque quality and addressing inflammatory and structural features of established plaques that lipid lowering alone may not fully resolve.

## ACKNOWLEDGEMENTS

This work was supported in part by the NIH award R56HL173828 (to K.-C. W.) and the Arizona Biomedical Research Centre grant RFGA2024-022-029 (to K.-C. W.). The authors acknowledge the use of the Regenerative Medicine Core and Eyring Materials Center. Schematics created using BioRender.com.

## AUTHOR CONTRIBUTIONS

Conceptualization: T.-Y.W. and K.-C.W. Methodology: T.-Y.W., A.Z., S.-P.W., H.-C. H., and C.L.P. Investigation: T.-Y.W., A.Z., S.-P.W., H.-C. H., and C.L.P. Visualization: T.-Y.W., A.Z., C.L.P., and K.-C.W. Supervision: K.-C.W. Writing –Original Draft: T.-Y.W. and K.-C.W.; Editing: T.Y.W., C.L.P., and K.-C.W.

## DATA AND MATERIALS AVAILABILITY

All data needed to evaluate the conclusions in the paper are present in the paper and/or the Supplementary Materials.

## REFERENCES

1. Jebari-Benslaiman S, Galicia-García U, Larrea-Sebal A, et al. Pathophysiology of Atherosclerosis. IJMS. 2022;23(6):3346. doi:10.3390/ijms23063346

2. Frostegård J. Immunity, atherosclerosis and cardiovascular disease. BMC Med. 2013;11(1):117. doi:10.1186/1741-7015-11-117

3. Ridker PM, Everett BM, Thuren T, et al. Antiinflammatory Therapy with Canakinumab for Atherosclerotic Disease. N Engl J Med. 2017;377(12):1119–1131. doi:10.1056/NEJMoa1707914

4. Tardif JC, Kouz S, Waters DD, et al. Efficacy and Safety of Low-Dose Colchicine after Myocardial Infarction. N Engl J Med. 2019;381(26):2497–2505. doi:10.1056/NEJMoa1912388

5. Hou P, Fang J, Liu Z, et al. Macrophage polarization and metabolism in atherosclerosis. Cell Death Dis. 2023;14(10):691. doi:10.1038/s41419-023-06206-z

6. Yang B, Hang S, Xu S, et al. Macrophage polarisation and inflammatory mechanisms in atherosclerosis: Implications for prevention and treatment. Heliyon. 2024;10(11):e32073. doi:10.1016/j.heliyon.2024.e32073

7. Wang KC, Yeh YT, Nguyen P, et al. Flow-dependent YAP/TAZ activities regulate endothelial phenotypes and atherosclerosis. Proc Natl Acad Sci USA. 2016;113(41):11525–11530. doi:10.1073/pnas.1613121113

8. Wang L, Luo JY, Li B, et al. Integrin-YAP/TAZ-JNK cascade mediates atheroprotective effect of unidirectional shear flow. Nature. 2016;540(7634):579–582. doi:10.1038/nature20602

9. Mao J, Yang R, Yuan P, et al. Different stimuli induce endothelial dysfunction and promote atherosclerosis through the Piezo1/YAP signaling axis. Archives of Biochemistry and Biophysics. 2023;747:109755. doi:10.1016/j.abb.2023.109755

10. Zhang X, Sun X, Qin Q, et al. YAP-mediated macrophage polarization is involved in progression of atherosclerosis. European Journal of Pharmacology. 2025;1008:178328. doi:10.1016/j.ejphar.2025.178328

11. Liu M, Yan M, Lv H, et al. Macrophage K63-Linked Ubiquitination of YAP Promotes Its Nuclear Localization and Exacerbates Atherosclerosis. Cell Reports. 2020;32(5):107990. doi:10.1016/j.celrep.2020.107990

12. Kimura TE, Duggirala A, Smith MC, et al. The Hippo pathway mediates inhibition of vascular smooth muscle cell proliferation by cAMP. Journal of Molecular and Cellular Cardiology. 2016;90:1–10. doi:10.1016/j.yjmcc.2015.11.024

13. Xiao J, Jin K, Wang J, et al. Conditional knockout of TFPI-1 in VSMCs of mice accelerates atherosclerosis by enhancing AMOT/YAP pathway. International Journal of Cardiology. 2017;228:605–614. doi:10.1016/j.ijcard.2016.11.195

14. Huang HC, Wang TY, Rousseau J, et al. Biomimetic nanodrug targets inflammation and suppresses YAP/TAZ to ameliorate atherosclerosis. Biomaterials. 2024;306:122505. doi:10.1016/j.biomaterials.2024.122505

15. Rousseau J, Wang T, McClendon S, et al. Monocyte-Mimetic Contrast Agent Enables Targeted and Sensitive Magnetic Resonance Imaging of Atherosclerotic Lesions. Adv Healthcare Materials. 2026;15(3):e02001. doi:10.1002/adhm.202502001

16. Nam D, Ni CW, Rezvan A, et al. Partial carotid ligation is a model of acutely induced disturbed flow, leading to rapid endothelial dysfunction and atherosclerosis. American Journal of Physiology-Heart and Circulatory Physiology. 2009;297(4):H1535–H1543. doi:10.1152/ajpheart.00510.2009

17. Adjei-Sowah EA, O’Connor SA, Veldhuizen J, et al. Investigating the Interactions of Glioma Stem Cells in the Perivascular Niche at Single-Cell Resolution using a Microfluidic Tumor Microenvironment Model. Adv Sci (Weinh). 2022;9(21):e2201436. doi:10.1002/advs.202201436

18. Lechner KS, Neurath MF, Weigmann B. Role of the IL-2 inducible tyrosine kinase ITK and its inhibitors in disease pathogenesis. J Mol Med (Berl). 2020;98(10):1385–1395. doi:10.1007/s00109-020-01958-z

19. Yang H, Zhou T, Stranz A, DeRoo E, Liu B. Single-Cell RNA Sequencing Reveals Heterogeneity of Vascular Cells in Early Stage Murine Abdominal Aortic Aneurysm-Brief Report. Arterioscler Thromb Vasc Biol. 2021;41(3):1158–1166. doi:10.1161/ATVBAHA.120.315607

20. Li F, Yan K, Wu L, et al. Single-cell RNA-seq reveals cellular heterogeneity of mouse carotid artery under disturbed flow. Cell Death Discov. 2021;7(1):180. doi:10.1038/s41420-021-00567-0

21. Müller AM, Hermanns MI, Skrzynski C, Nesslinger M, Müller KM, Kirkpatrick CJ. Expression of the Endothelial Markers PECAM-1, vWf, and CD34 in Vivo and in Vitro. Experimental and Molecular Pathology. 2002;72(3):221–229. doi:10.1006/exmp.2002.2424

22. Andreatta M, Corria-Osorio J, Müller S, Cubas R, Coukos G, Carmona SJ. Interpretation of T cell states from single-cell transcriptomics data using reference atlases. Nat Commun. 2021;12(1):2965. doi:10.1038/s41467-021-23324-4

23. Liu T, Chen Y, Hou L, et al. Immune cell-mediated features of atherosclerosis. Front Cardiovasc Med. 2024;11:1450737. doi:10.3389/fcvm.2024.1450737

24. Liu X, Li X, Wang X, et al. Single-cell RNA-seq analysis of mouse carotid artery under disturbed flow and human carotid plaques identifies key cell populations in atherosclerosis development. Sci Rep. 2025;15(1):20747. doi:10.1038/s41598-025-07395-7

25. Zernecke A, Erhard F, Weinberger T, et al. Integrated single-cell analysis-based classification of vascular mononuclear phagocytes in mouse and human atherosclerosis. Cardiovascular Research. 2023;119(8):1676–1689. doi:10.1093/cvr/cvac161

26. Peled M, Nishi H, Weinstock A, et al. A wild-type mouse-based model for the regression of inflammation in atherosclerosis. Schulz C, ed. PLoS ONE. 2017;12(3):e0173975. doi:10.1371/journal.pone.0173975

27. Goncalves I, Pan M, Singh P, et al. Spatial transcriptomics reveals a key role of fibroblast-like vascular smooth muscle cells in human atherosclerotic cell crosstalk and stability. European Heart Journal. Published online February 13, 2026:ehaf1091. doi:10.1093/eurheartj/ehaf1091

28. Cipollone F, Fazia M, Mincione G, et al. Increased Expression of Transforming Growth Factor-β1 as a Stabilizing Factor in Human Atherosclerotic Plaques. Stroke. 2004;35(10):2253–2257. doi:10.1161/01.STR.0000140739.45472.9c

29. Mallat Z, Gojova A, Marchiol-Fournigault C, et al. Inhibition of Transforming Growth Factor-β Signaling Accelerates Atherosclerosis and Induces an Unstable Plaque Phenotype in Mice. Circulation Research. 2001;89(10):930–934. doi:10.1161/hh2201.099415

30. Edsfeldt A, Singh P, Matthes F, et al. Transforming growth factor-β2 is associated with atherosclerotic plaque stability and lower risk for cardiovascular events. Cardiovascular Research. 2023;119(11):2061–2073. doi:10.1093/cvr/cvad079

31. Roche-Molina M, Sanz-Rosa D, Cruz FM, et al. Induction of Sustained Hypercholesterolemia by Single Adeno-Associated Virus–Mediated Gene Transfer of Mutant hPCSK9. ATVB. 2015;35(1):50–59. doi:10.1161/ATVBAHA.114.303617

32. Keeter WC, Carter NM, Nadler JL, Galkina EV. The AAV-PCSK9 murine model of atherosclerosis and metabolic dysfunction. Ketelhuth D, ed. European Heart Journal Open. 2022;2(3):oeac028. doi:10.1093/ehjopen/oeac028

33. Bjørklund MM, Hollensen AK, Hagensen MK, et al. Induction of Atherosclerosis in Mice and Hamsters Without Germline Genetic Engineering. Circulation Research. 2014;114(11):1684–1689. doi:10.1161/CIRCRESAHA.114.302937

34. Zhang HL, Wu J, Zhu J. The Immune-Modulatory Role of Apolipoprotein E with Emphasis on Multiple Sclerosis and Experimental Autoimmune Encephalomyelitis. Blaser K, ed. Journal of Immunology Research. 2010;2010(1):186813. doi:10.1155/2010/186813

35. Hansson GK, Libby P, Tabas I. Inflammation and plaque vulnerability. J Intern Med. 2015;278(5):483–493. doi:10.1111/joim.12406

36. Spagnoli LG, Bonanno E, Sangiorgi G, Mauriello A. Role of Inflammation in Atherosclerosis. Journal of Nuclear Medicine. 2007;48(11):1800–1815. doi:10.2967/jnumed.107.038661

37. Hoeffner N, Paul A, Goo YH. Drug screen identifies verteporfin as a regulator of lipid metabolism in macrophage foam cells. Sci Rep. 2023;13(1):19588. doi:10.1038/s41598-023-46467-4

38. Zhang X, Sun X, Qin Q, et al. YAP-mediated macrophage polarization is involved in progression of atherosclerosis. Eur J Pharmacol. 2025;1008:178328. doi:10.1016/j.ejphar.2025.178328

39. Pan H, Xue C, Auerbach BJ, et al. Single-Cell Genomics Reveals a Novel Cell State During Smooth Muscle Cell Phenotypic Switching and Potential Therapeutic Targets for Atherosclerosis in Mouse and Human. Circulation. 2020;142(21):2060–2075. doi:10.1161/CIRCULATIONAHA.120.048378

40. Winkels H, Ehinger E, Vassallo M, et al. Atlas of the Immune Cell Repertoire in Mouse Atherosclerosis Defined by Single-Cell RNA-Sequencing and Mass Cytometry. Circ Res. 2018;122(12):1675–1688. doi:10.1161/CIRCRESAHA.117.312513

41. Wirka RC, Wagh D, Paik DT, et al. Atheroprotective roles of smooth muscle cell phenotypic modulation and the TCF21 disease gene as revealed by single-cell analysis. Nat Med. 2019;25(8):1280–1289. doi:10.1038/s41591-019-0512-5

42. Yurdagul A. Crosstalk Between Macrophages and Vascular Smooth Muscle Cells in Atherosclerotic Plaque Stability. Arterioscler Thromb Vasc Biol. 2022;42(4):372–380. doi:10.1161/ATVBAHA.121.316233

43. Newby AC. Metalloproteinase expression in monocytes and macrophages and its relationship to atherosclerotic plaque instability. Arterioscler Thromb Vasc Biol. 2008;28(12):2108–2114. doi:10.1161/ATVBAHA.108.173898

44. Bi Y, Chen J, Hu F, Liu J, Li M, Zhao L. M2 Macrophages as a Potential Target for Antiatherosclerosis Treatment. Neural Plast. 2019;2019:6724903. doi:10.1155/2019/6724903

45. Zhang H, Ramakrishnan SK, Triner D, et al. Tumor-selective proteotoxicity of verteporfin inhibits colon cancer progression independently of YAP1. Sci Signal. 2015;8(397):ra98. doi:10.1126/scisignal.aac5418

46. Zhou W, Lim A, Elmadbouh OHM, et al. Verteporfin induces lipid peroxidation and ferroptosis in pancreatic cancer cells. Free Radic Biol Med. 2024;212:493–504. doi:10.1016/j.freeradbiomed.2024.01.003

47. Wei H, Wang F, Wang Y, et al. Verteporfin suppresses cell survival, angiogenesis and vasculogenic mimicry of pancreatic ductal adenocarcinoma via disrupting the YAP-TEAD complex. Cancer Sci. 2017;108(3):478–487. doi:10.1111/cas.13138

48. Shah SR, Kim J, Schiapparelli P, et al. Verteporfin-Loaded Polymeric Microparticles for Intratumoral Treatment of Brain Cancer. Mol Pharm. 2019;16(4):1433–1443. doi:10.1021/acs.molpharmaceut.8b00959

49. Golino JL, Wang X, Bian J, et al. Anti-Cancer Activity of Verteporfin in Cholangiocarcinoma. Cancers (Basel). 2023;15(9):2454. doi:10.3390/cancers15092454

