## Supplementary information for "Monocyte-Mimetic Nanoparticle Delivery of Verteporfin Promotes Plaque-Stabilizing Remodeling in Atherosclerosis"

**TITLE**

**AFFILIATION**

**Supplementary Figure 1**

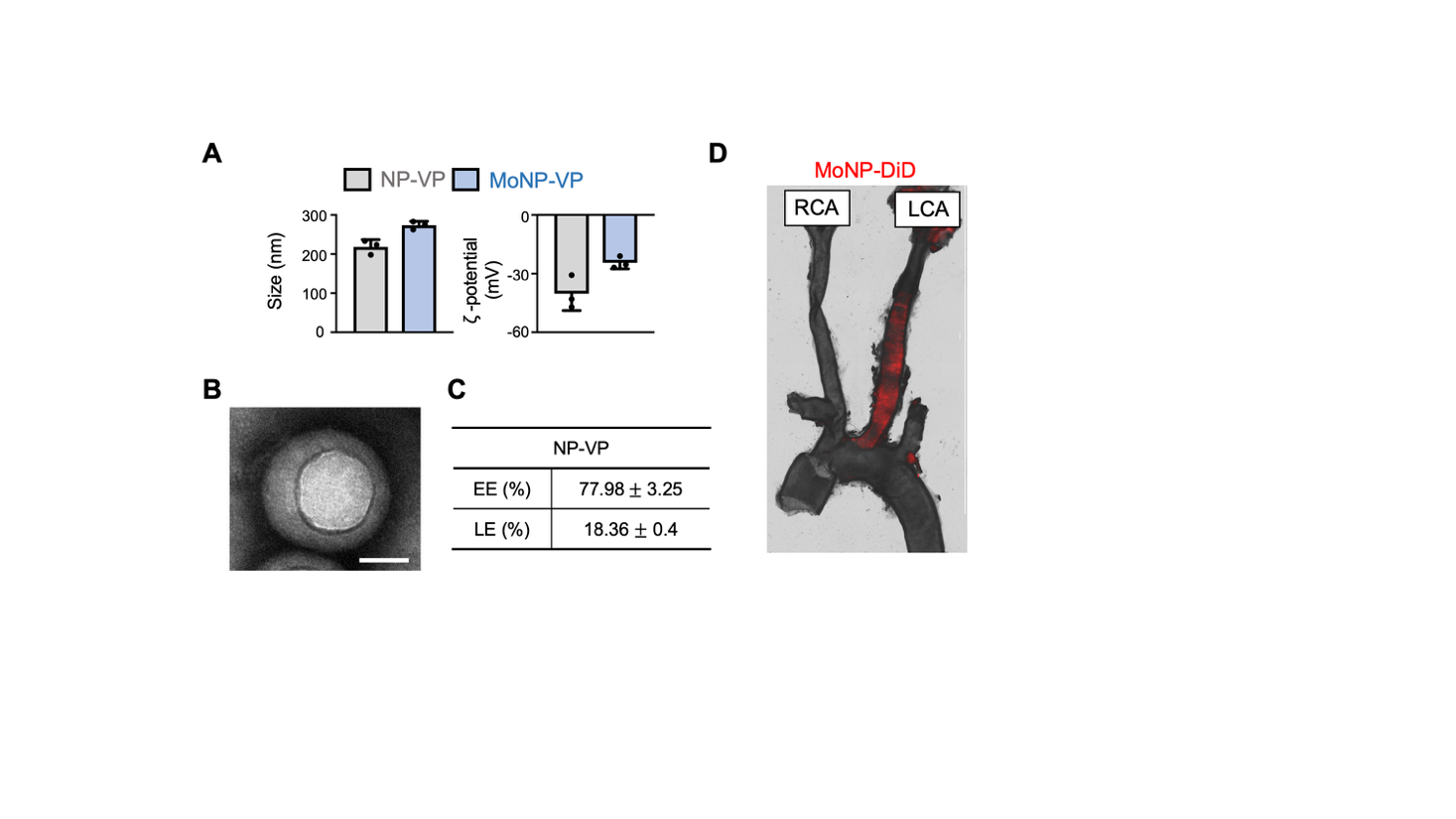

**Supplementary Figure 1. Characterization of monocyte coated nanoparticles (MoNP).** (A) Hydrodynamic size and zeta potential of MoNP-VP and MoNP. (B) TEM image of MoNP-VP. Scale bar = 50 nm. (C) Encapsulation efficiency (EE) and loading efficiency (LE) of VP in MoNP-VP. (D) Representative image showing MoNP-DiD accumulation in the partially ligated carotid artery. Data are presented as mean ± SD. n=3.

**Supplementary Figure 2**

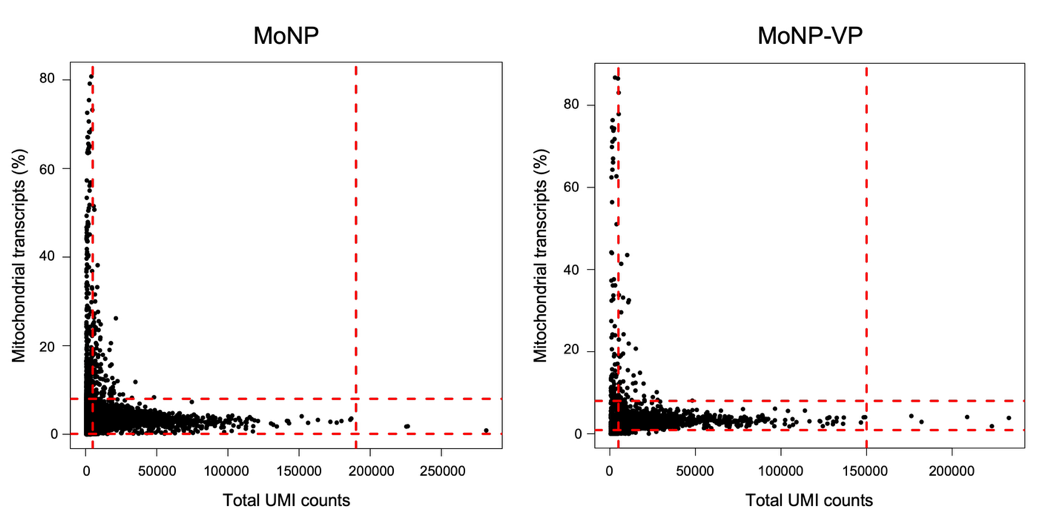

**Supplementary Figure 2. Quality control analysis of scRNA-seq data.** Scatter plots showing mitochondrial transcript percentage versus total UMI counts for cells isolated from MoNP and MoNP-VP treated arteries. Red dashed lines indicate the filtering thresholds used for quality control analysis.

**Supplementary Figure 3**

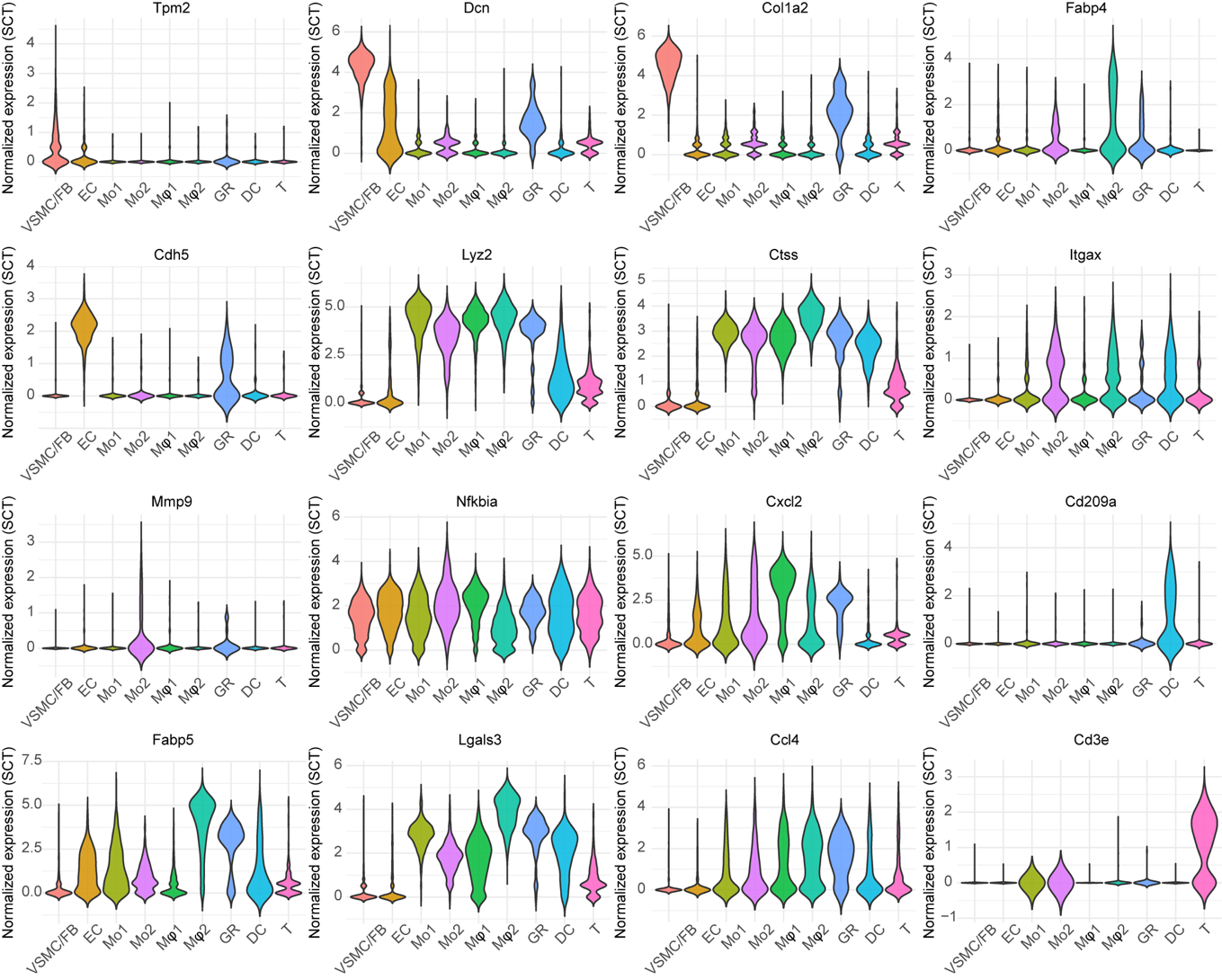

**Supplementary Figure 3. Expression of additional marker genes across annotated cell populations.** Violin plots showing the normalized expression of selected marker genes across annotated cell populations identified by scRNA-seq.

**Supplementary Figure 4**

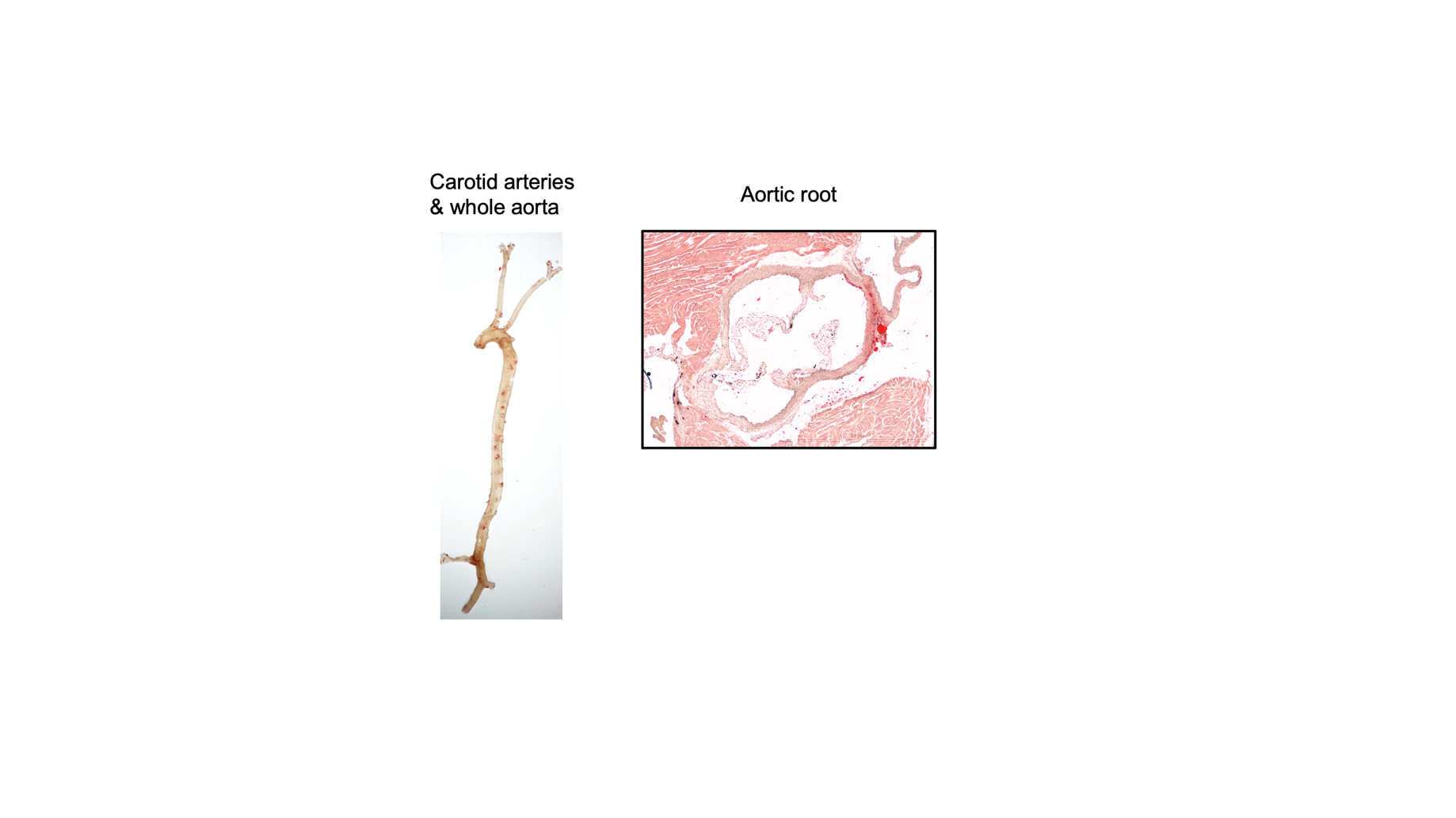

**Supplementary Figure 4. Atherosclerotic lesion formation in saline-injected mice.** Representative Oil red O image of carotid arteries, whole aortas, and aortic root sections from saline-injected mice. n=3

**Supplementary Table 1: Serum metabolic panel analysis**

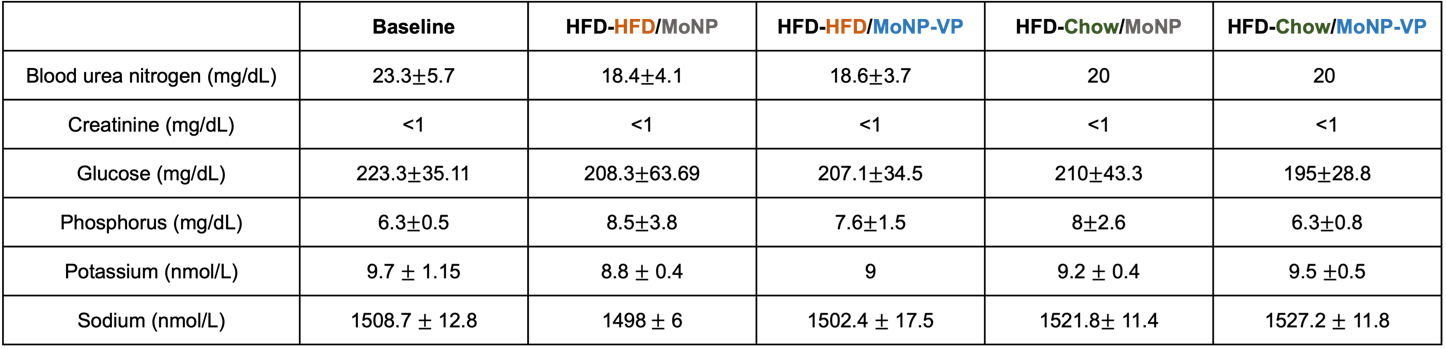

**Supplementary Table 2: Primer list**

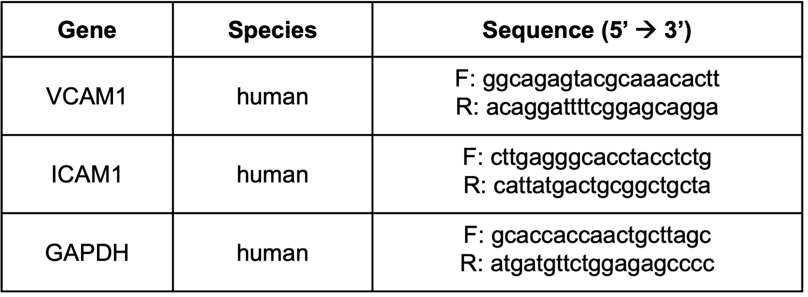
